# When GWAS meets the Connectivity Map: drug repositioning for seven psychiatric disorders

**DOI:** 10.1101/096503

**Authors:** Hon-Cheong So, Carlos K.L. Chau, Wan-To Chiu, Kin-Sang Ho, Cho-Pong Lo, Stephanie Ho-Yue Yim, Pak C. Sham

## Abstract

Our knowledge of disease genetics has advanced rapidly during the past decade, with the advent of high-throughput genotyping technologies such as genome-wide association studies (GWAS). However, few methodologies were developed and systemic studies performed to identify novel drug candidates utilizing GWAS data. In this study we focus on drug repositioning, which is a cost-effective approach to shorten the developmental process of new therapies. We proposed a novel framework of drug repositioning by comparing GWAS-imputed transcriptome with drug expression profiles from the Connectivity Map. The approach was applied to 7 psychiatric disorders. We discovered a number of novel repositioning candidates, many of which are supported by preclinical or clinical evidence. We found that the predicted drugs are significantly enriched for known psychiatric medications, or therapies considered in clinical trials. For example, drugs repurposed for schizophrenia are strongly enriched for antipsychotics (*p* = 4.69E-06), while those repurposed for bipolar disorder are enriched for antipsychotics (*p* = 2.26E-07) and antidepressants (*p* = 1.17E-05). These findings provide support to the usefulness of GWAS signals in guiding drug discoveries and the validity of our approach in drug repositioning. We also present manually curated lists of top repositioning candidates for each disorder, which we believe will serve as a useful resource for researchers.

## INTRODUCTION

The last decade has witnessed a rapid growth of genotyping technologies and genome-wide association studies (GWAS) have played in major role in unraveling the genetic bases of complex diseases. According to the latest GWAS catalog (www.ebi.ac.uk/gwas. accessed 22^nd^ Dec 2016), more than 2600 GWAS studies have been performed to date. It is worth noting that many biobanks are also collecting genomic data. and the private genetics company 23andMe has genotyped more than a million customers (mediacenter.23andme.com/fact-sheet/). It is clearly of great clinical and public interest to translate these findings into treatment for diseases. Nevertheless, compared to the large literature of GWAS studies, relatively few methodologies have been developed and systematic studies performed to identify novel drug candidates by using GWAS data.

Psychiatric disorders carry significant burden on health globally^1^ and the current treatment strategies are far from perfect. Despite the heavy health burden and the increased awareness of mental health in many places, drug discovery in the field have largely been stagnant^2^. As argued by Hyman, the basic mechanisms of most antidepressants and antipsychotics, the most widely used drugs used in psychiatry, are relatively similar to their prototypes imipramine and chlorpromazine discovered in the 1950s^2^. On the other hand, in recent years GWAS studies have greatly advanced our knowledge of the genetic bases of many psychiatric disorders. Taking advantage of these developments, we proposed a new framework for identifying drug candidates based on GWAS results, and applied the method to a variety of psychiatric disorders.

Here we focus on drug repositioning, that is, finding new indications for existing drugs. As conventional drug development is an expensive and lengthy process, repositioning serves as a useful strategy to hasten the development cycle^3^. It is worth noting that while we use the term “drug repositioning” throughout this paper, the method is also applicable to any chemicals with known gene expression profiles.

A few previous studies have investigated the use of GWAS data in drug repositioning. The most intuitive approach is to study whether the top genes identified in GWAS can serve as drug targets. Sanseau et al.^4^ searched for top susceptibility genes (with *p* < 1e-7) from the GWAS catalog and matched the results against the drug targets listed in the Pharmaprojects database, a commercial resource with information on global drug pipelines. In addition, they proposed a repositioning approach by identifying “mismatches” between drug indications and the original GWAS traits. In another study^5^, Cao et al. considered interacting partners of GWAS hits to discover new drug targets. Lencz and Malhotra^6^ compared schizophrenia GWAS results with drug target genes, and prioritized potential new drug targets for the disease. Another recent study investigated whether evidence from human genetic studies are useful to drug development in general. The authors found that the proportion of drugs with support from GWAS increased along the development pipeline^7^.

While the approach of finding overlap between top GWAS hits and known drug target genes is useful, it has a number of limitations. Firstly, many of the top GWAS genes may not be easily targeted by a drug. Cao and Moult^5^ studied 856 drug target genes from DrugBank and found only 20 genes that were discovered in GWAS of the corresponding disease. While this number is likely to improve with increasing sample sizes, there will remain a number of GWAS hits which are not directly “druggable”. In addition, many of the GWAS top SNPs are within non-coding regions^8^ and do not encode for a drug target protein. The above approach might also miss “multi-target” drugs, a paradigm that has attracted increasing attention in recent years. It is argued that as complex diseases (like most psychiatric disorders) involve the interplay of multiple genetic and environmental factors, they may be more easily managed by modulating multiple instead of single targets^9^. Lastly, as previous studies mainly focused mainly on the most significant hits, they ignored the contribution of genetic variants with smaller effect sizes. As shown in polygenic score analyses of many complex traits, often variants achieving lower significance levels also contribute to disease risks^10^.

With the aforementioned limitations in mind, we developed a new strategy for drug repositioning by imputing gene expression profile from GWAS summary statistics and comparing it with drug-induced expression changes. Analysis of the transcriptome of drugs versus diseases is an established approach for drug repositioning and has previously been successfully applied for complex diseases^11,12^. For example, by examining drugs in the Connectivity Map (Cmap)^13^ which showed opposite patterns of expression to diseases, Sirota et al.^11^ derived candidates for repositioning and experimentally validated a prediction cimetidine for the treatment of lung adenocarcinoma. With a similar method, Dudley et al.^12^ identified topiramate as a novel treatment for inflammatory bowel disease and validated it in an animal model. Other studies (*e.g.*^14–16^) also showed potential of this approach in repositioning.

Built on this repositioning strategy, we proposed a new approach by using imputed transcriptome from GWAS instead of expression data from microarray or RNA-sequencing studies. This approach has several advantages. Firstly, patients from expression studies are often medicated. This is particularly relevant for studies in neuro-psychiatry, as brain tissues can only come from post-mortem samples of patients, who were often on psychiatric medications^17^. History of medications might confound our results as our aim is to compare the expression patterns of disease and those of drugs. Imputed transcriptome on the other hand is not altered by medications or other environmental confounders. Secondly, current GWAS samples are usually orders of magnitude larger than expression studies (often 10^4^ or more), and GWAS summary statistics are widely available. In addition, for many diseases such as psychiatric disorders, the tissues of interest are not easily accessible. On the other hand, as we shall explain below, expression profiles can be readily imputed for over 90 tissues from GWAS data using appropriate statistical models.

## METHODS

### Imputation of expression profile from GWAS data

Recently methods have been developed to impute expression from GWAS variants^18–20^. The main idea is to build a statistical model to predict expression levels from SNPs in a reference transcriptome dataset, and the prediction model can be applied to new genotype data. This approach estimates the component of gene expression that is contributed by (germline) genetic variations. The program PrediXcan^18^ was developed for this purpose based on models built from the Genotype-Tissue Expression (GTEx) project^21^ and the Depression Genes and Networks (DGN) cohort^22^. As most individual genotype data are not publicly available due to privacy concerns, we applied a recently developed algorithm called MetaXcan which allows imputation of expression *z*-scores (i.e. z-statistic derived from comparing gene expression in cases versus controls) based on summary statistics alone. The method was shown to be in excellent concordance with predictions made from raw genotype data^19^. Assuming that a set of SNPs (SNP_1_, SNP_2_ …, SNP_*k*_) contribute to the expression of gene g, the following formula can be used to compute the expression *z*-scores for the gene:

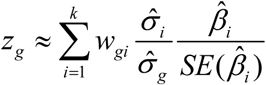

where *w_gi_* is the weight given to SNP_*i*_ for predicting the expression level of gene g, 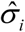 and 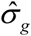 denote the estimated variance of SNP_*i*_; and gene *g* respectively (estimated from a reference genotype dataset), and 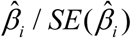 denotes the summary z-statistic of SNP_*i*_; of the disease trait. The weights of SNPs *w_gi_* were computed from reference datasets of expression quantitative trait loci (eQTL) studies. We downloaded pre-computed weights derived from an elastic net prediction model with GTEx as the reference transcriptome dataset, provided by the authors of MetaXcan. It is worth noting that the above methodology is similar to another transcriptome imputation algorithm TWAS^20^, but a different prediction method (based on a mixed model) is employed in TWAS. We used MetaXcan in this study as the program readily allows transcriptome imputation for a much larger variety of tissues.

### Transcriptome imputation of 7 psychiatric disorders

GWAS summary statistics were obtained from the Psychiatric Genomics Consortium (PGC) website (www.med.unc.edu/pgc/results-and-downloads). We downloaded eight sets of GWAS summary statistics corresponding to seven psychiatric disorders, including schizophrenia (SCZ)^23^, major depressive disorder (MDD)^24,25^, bipolar disorder (BD)^26^, Alzheimer’s disease (AD)^27^, anxiety disorders (ANX)^28^, autistic spectrum disorders (ASD)^29^ and attention deficit hyperactivity disorder (ADHD)^30^. We employed two different sets of summary statistics for MDD; the first set was from the PGC group^24^, while the other was from the CONVERGE study which recruited more severe MDD cases from Chinese women only^25^. Details of individual studies are described in the respective references. Transcriptome of each disease was imputed for ten brain regions included in GTEx, using default parameters in MetaXcan.

### Drug-induced expression profiles

Drug-induced expression profiles were derived from the Cmap database, a resource of genome-wide expression profile of cultured cell lines treated with 1309 different drugs or small molecules^13^. We downloaded raw expression data from Cmap, and performed normalization with the MAS5 algorithm. Expression levels of genes represented on more than one probe sets were averaged. Differential expression between treated cell lines and controls was tested using the limma package^31^. We performed analyses on each combinations of drug and cell line, with a total of 3478 instances. Statistical analyses were performed in R3.2.1 with the assistance of the R package “longevityTools”.

### Comparison of gene expression profiles of drugs versus diseases

Next we compared the expression profiles (in z-scores) of drugs versus those of diseases. The original study on Cmap employed Kolmogorov-Smirnov (KS) tests^13^ to compare two expression patterns. In brief, the aim of the KS test is to evaluate whether a set of disease-related genes are ranked higher or lower than expected in a list of genes sorted by their drug-induced expression levels. The KS test was performed separately for upregulated and downregulated disease genes. For drug repositioning, we study whether there is an enrichment of genes that are upregulated for disease but downregulated on drug treatment, and vice versa. We adopted the same formulae described in original Cmap paper to calculate the “connectivity scores”.

Reversed patterns of expression can also be tested by Spearman or Pearson correlations. Yet another approach is to use only the *K* most up- or down-regulated genes in computing the correlations^32^. In this study we employed all five methods (*i.e.* KS test, Spearman correlation with all or the most differentially expressed genes, Pearson correlation with all or the most differentially expressed genes) in our analyses and computed the average ranks. As there are no consensus methods to define *K*, we also set different values of *K* (50, 100, 250, 500) and averaged the results for each method. Drugs were then ranked in ascending order of their connectivity scores or correlation results (*i.e.* the most negative correlation or connectivity score ranked first).

To assess the significance of the ranks, we performed permutation tests by shuffling the disease expression z-scores and comparing it to drug transcriptomic profiles. One hundred permutations were performed for each drug-disease pair, and the distribution of ranks under the null was combined across all drug-disease pairs (such that the null distribution was derived from 347800 ranks under H_0_).

### Manual curations of the top repositioning candidates

To assess the drug candidates found in our drug repositioning algorithm, we performed manual curations of the top 15 drugs (representing the top ~0.45% of all instances) identified for each disorder and brain region. Literature search was performed to look for evidence of therapeutic potential of the identified drug candidates.

### Tests for enrichment of known indicated drugs or drugs in clinical trial

In addition to manually inspecting the top candidates, in order to validate our approach, we also tested for an enrichment of drugs that are (1) indicated for each disorder; or (2) included in clinical trials. The enrichment tests are similar in principle to gene-set analyses, but with gene-sets replaced by drug-sets. We employed two types of tests for enrichment. In the first approach, we tested whether a known drug-set, such as antipsychotics or antidepressants, were ranked significantly higher than by chance; this approach is also known as a “self-contained” test. In the second method, we compared medications in the drug-set against those outside the set, and tested whether the former group was ranked significantly higher. This is also known as a “competitive” test^33,34^. Details of the statistical methods are described in Supplementary Text.

We considered three sources of drug-sets in our analyses. The first set comes from the Anatomical Therapeutic Classification (ATC) drugs downloaded from KEGG. The ATC is an established system for the classification of medications and we extracted three groups of drugs: (1) all psychiatric drugs (coded “N05” or “N06”); (2) antipsychotics (coded “N05A”); (3) antidepressants and anxiolytics (coded “N05B” or “N06A”). We grouped antidepressants and anxiolytics together in our analyses as many anti-depressants are indicated for anxiety disorders, and vice versa, anxiolytics are frequently prescribed to depressive patients clinically^35^. We did not specifically include drugs for dementia or psychostimulants as they are relatively few in number. Note that the ATC does not classify drugs specifically for some disorders, such as BD, autism and ADHD.

The second source is from Wei et al.^36^ who complied a MEDication Indication resource (MEDI) from four public medication resources, including RxNorm, Side Effect Resource 2 (SIDER2), Wikipedia and MedlinePlus. A random subset of the extracted indications was also reviewed by physicians. We used the MEDI high-precision subset (MEDI-HPS) which only include drug indications found in RxNorm or in at least 2 out of 3 other sources, with an estimated precision of 92%.

As the aforementioned sources only include known drug indications, we also considered a wider set of drugs that are included for clinical trials on https://clinicaltrials.gov. These drugs represent promising candidates that are often supported by preclinical or human studies. We downloaded a precompiled list of these drugs (created in May 2016) from https://doi.org/10.15363/thinklab.d212.

For each disorder, we compiled a list of repositioning candidates using the imputed transcriptome profile of each brain region. We combined the drug-set analysis results across brain regions by meta-analyses of the respective p-values. We employed two different algorithms, Fisher’s method^37^ and Tippett’s minimum *p* approach^38^ to combine the p-values. Analyses were performed with the R package “metap”. Multiple testing correction was performed by the false discovery rate (FDR) approach.

## RESULTS

The sample sizes of the GWAS datasets we used are listed in Supplementary Table 1. The top 15 repositioned drug candidates for each disorder and brain region (with manual curations of drug descriptions and potential therapeutic relationship to the disorder) are presented in full in Supplementary Tables 2–9. Selected drug candidates within the top lists are presented and discussed below.

**Table 1.** Selected drug repositioning candidates for schizophrenia

| Drug | Cell line | Brain<br>Region | Rank | *p*-value | Brief Description |
| --- | --- | --- | --- | --- | --- |
| Tremorine | PC3 | AC | 7 | 3.19E-03 | Affect amine levels in brain; a closely related drug oxotremorine was effective for SCZ in animal studies |
| Scopolamine | HL60 | CAU | 3 | 1.43E-03 | Muscarinic antagonist; antidepressant properties |
| Ranitidine | MCF7 | CAU | 12 | 3.36E-03 | H2 antagonist; shown to reduce negative symptoms in clinical study |
| Paroxetine | PC3 | CAU | 14 | 4.59E-03 | SSRI; shown to improve negative symptoms in clinical study |
| Droperidol | HL60 | CER | 4 | 5.12E-04 | D2 antagonist; antipsychotic properties |
| Thiopropazine | PC3 | CEH | 7 | 2.56E-03 | First-generation anti-psychotic |
| Tetrahydroalstonine | PC3 | COR | 1 | 7.19E-05 | Indole alkaloid; anti-psychotic properties in rodents; clinically used in Nigeria |
| Hesperidin | MCF7 | COR | 2 | 8.91E-05 | Flavanones ; antioxidant |
| Trifluoperazine | PC3 | COR | 5 | 8.40E-04 | First-generation anti-psychotic |
| Bumetanide | MCF7 | COR | 6 | 8.57E-04 | Na-K-Cl cotransporter 1 inhibitor; improved hallucinations in RCT |
| Meclofenamic acid | MCF7 | FCOR | 9 | 2.24E-03 | NSAID; under clinical trial for cognitive symptoms |
| Spiradoline | PC3 | FCOR | 10 | 2.24E-03 | Selective kappa-Opioid Agonist; effective in animal studies |
| Thiethylperazine | PC3 | FCOR | 13 | 3.40E-03 | First-generation anti-psychotic |
| Idazoxan | PC3 | FOCR | 14 | 3.64E-03 | Alpha 2 antagonist; clinical studies showed potential in improving negative symptoms |
| Bromopride | MCF7 | HYP | 4 | 1.46E-03 | Dopamine antagonist |
| Spiperone | PC3 | NUC | 8 | 3.37E-03 | First-generation antipsychotic |
| Pimozide | PC3 | NUC | 13 | 4.14E-03 | First-generation antipsychotic |
| Metitepine | PC3 | PUT | 1 | 5.75E-06 | 5-HT and dopamine receptor antagonist with possible antipsychotic properties |

**Table 2.** Selected drug repositioning candidates for bipolar disorder

| Drug | Cell line | Brain<br>Region | Rank | *p*-value | Brief Description |
| --- | --- | --- | --- | --- | --- |
| Norcycllobenzaprine | PC3 | CAU | 1 | 3.31E-04 | Structurally very similar to the antidepressant amitriptyline |
| Acetylsalicylsalicylic acid | PC3 | CAU | 2 | 3.65E-04 | Closely related to aspirin, which may be useful for BD |
| Thioridazine | HL60 | CAU | 6 | 1.03E-03 | First generation antipsychotic |
| Acetylsalicylic acid | HL60 | CEH | 9 | 2.00E-03 | Aspirin; clinical studies showed potential in BD treatment |
| Mexiletine | PC3 | CER | 6 | 2.25E-03 | Class IB anti-arrhythmic; may be useful for treatment resistant BD |
| Dextromethorphan | HL60 | CER | 11 | 4.23E-03 | Morphinan class; may be effective for bipolar depression |
| Imipramine | HL60 | CER | 14 | 5.56E-03 | Tricyclic antidepressant |
| Simvastatin | MCF7 | COR | 6 | 5.59E-03 | Augmentation with lithium may be effective for BD |
| Metirapone | HL60 | FCOR | 1 | 1.58E-04 | Cortisol synthesis inhibitor; RCT showed effects in treatment-resistant MDD |
| Prestwick-689<br>(androsterone) | MCF7 | COR | 13 | 2.53E-03 | DHEA (precursor of this drug) may have anti-depressant effects |
| Alpha yohimbine | MCF7 | HIP | 1 | 3.82E-04 | Alpha 2 antagonist; an RCT showed yohimbine hastened anti-depressant response if combined with fluoxetine |
| Trifluoperazine | MCF7 | HIP | 5 | 1.22E-03 | Antipsychotic |
| Molindone | MCF7 | HIP | 12 | 3.72E-03 | Antipsychotic |
| Metformin | MCF7 | HIP | 13 | 3.90E-03 | Antidiabetic; being tested in clinical trial for refractory BD |
| SC.58125 | MCF7 | HYP | 4 | 1.12E-03 | COX-2 inhibitor; a closely related drug celecoxib showed antidepressant actions in BD |
| Ketoconazole | PC3 | HYP | 7 | 2.27E-03 | Case reports and open studies suggested efficacy for bipolar depression |
| Thiopropazine | MCF7 | HYP | 10 | 3.54E-03 | First generation antipsychotic |
| Mesoridazine | PC3 | NUC | 6 | 1.99E-03 | First generation antipsychotic |
| Metitepine | PC3 | PUT | 1 | 5.75E-06 | 5-HT and dopamine receptor antagonist with possible antipsychotic properties |

### Drug candidates repositioned for different psychiatric disorders

#### Schizophrenia

Table 1 shows selected candidates for schizophrenia and bipolar disorder. For schizophrenia, it is interesting to note that we identified a number of known antipsychotics such as thioproperazine, droperidol, triflupromazine, thiethylperazine, spiperone and pimozide as top candidates. It is worth noting that our repositioning method is blind to any knowledge about existing psychiatric drugs or known drug targets. The results provide further evidence to the role of the dopaminergic system in the treatment of schizophrenia. Some top candidates have been shown to improve negative symptoms in clinical studies, such as the H2 antagonist ranitidine^39^ and the alpha 2 receptor antagonist idazoxan^40^. The antidepressant paroxetine was also tested in a double-blind clinical trial and shown to be efficacious in ameliorating negative symptoms in schizophrenia^41^. There are other drug candidate with preliminary support by preclinical or clinical studies with diverse mechanisms, such as the serotonin and dopamine antagonist metitepine^42^, the Na-K-Cl cotransporter 1 inhibitor bumetanide^43^ and the non-steroidal anti-inflammatory drug (NSAID) meclofenamic acid^44^. It is also noteworthy that a relatively large proportion of drugs with stronger literature support were derived from comparison with expressions in the frontal cortex, a brain region strongly implicated in schizophrenia^45^.

#### Bipolar disorder

As for bipolar disorder, we found a number of antipsychotics among the top list (Table 2). Antipsychotics are well-known to be effective for BD overall and for the associated psychotic symptoms. Our analyses also revealed other candidates with known or potential antidepressant effects, such as imipramine, yohimbine, metyrapone^46^ and ketoconazole^47^. The latter two drugs are believed to exert antidepressant effects by reducing cortisol levels. Antidepressants are often used in the treatment of bipolar patients^48^. While there are controversies regarding its use, anti-depressants are included as valid treatment options in current guidelines, especially in bipolar II patients or when used with a mood stabilizer^49^. Interestingly, a few NSAIDs were also on the top list, such as aspirin and a cyclooxygenase-2 (COX-2) inhibitor SC-58125, which is supported by the neuro-inflammatory hypothesis of BD^50^. In line with the observation of raised cardiovascular risks in bipolar patients and possible shared pathophysiology between these disorders^51^, simvastatin and metformin were also among the top list.

#### Major depressive disorder

Two sets of GWAS data for MDD were used in our analyses (Table 3). For the results using the MDD-PGC data, we observed that fluoxetine, a widely used selective serotonin reuptake inhibitor (SSRI), was among the top candidates. We also observed quite a few antipsychotic medications on the list, such as sulpride, promazine, perphenazine and loxapine. Other repositioning candidates are more diverse in their mechanisms, such as phosphodiesterase inhibitors (papaverine^52^), histamine receptor antagonists (thioperamide^52^), muscarinic antagonists (scopolamine^53^) and cortisol-lowering agents (ketoconazole^47^) etc.

**Table 3.**
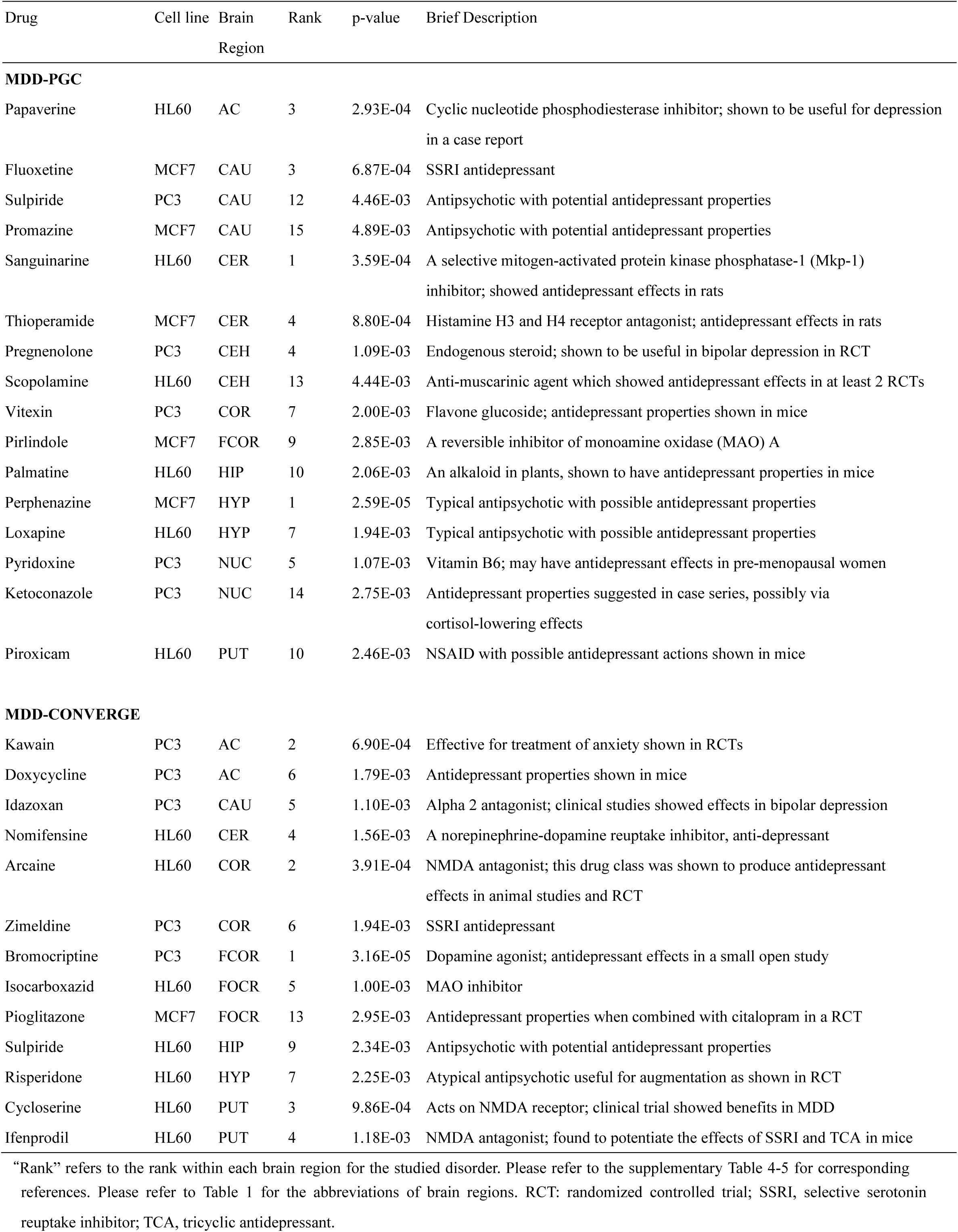
Selected drug repositioning candidates for major depressive disorder

As for the repositioning results from the MDD-CONVERGE study, a known SSRI (zimeldine^54^), a norepinephrine-dopamine reuptake inhibitor (nomifensine^55^) and a monoamine oxidase (MAO) inhibitor (isocarboxazid) were ranked among the top, although the former two drugs have been withdrawn due to other unrelated adverse effects. Two antipsychotics sulpride and risperidone were also on the list. Sulpride was tested in a double blind randomized controlled trial (RCT) of 177 patients and shown to significantly improve depressive symptoms^56^. Risperidone was confirmed to be useful as adjunctive treatment for MDD^57^. Interestingly, we found three drugs with actions on the glutamatergic system among the top candidates^58^. Arcaine and ifenprodil are NMDA antagonists and cycloserine is a partial agonist at the glycine site of the NMDA glutamate receptor. Ifenprodil was shown to improve depression in a mouse model^59^ while cycloserine has been shown to be effective as an add-on therapy in an RCT^60^.

#### Anxiety disorders

For anxiety disorders, we found the SSRI paroxetine and the tricyclic antidepressant protriptyline were on the top list. Another drug on the serotonergic system, pirenperone acts as a 5-HT2 antagonist and was shown anxiolytic actions in a small clinical study^61^. Some other drugs with preliminary support by animal or clinical studies include bumetanide, a loop diuretic which also affects GABA_A_ signaling^62^; ivermectin, an anti-parasitic and GABA agonist^63^; kawain, a kavalactone with anxiolytic properties shown in a number of clinical trials^64^; nimodipine, a calcium channel blocker with potential anxiolytic effects^65^ and piracetam, a nootropic drug which is also proposed for treatment of dementia^66^.

#### Alzheimer disease

AD is a relatively intense research area and we have found numerous drugs with some support by preclinical or clinical studies. Some of the drugs among the top candidates received have been tested in clinical trials. Naftidrofuryl is a vasodilatory agent which was shown to be effective for functional outcomes, mood and cognitive function in a meta-analysis^67^. Vinpocetine is an alkaloid from a plant which showed some benefits in dementia according to a meta-analysis of three RCTs^68^. Some other interesting repositioning candidates include verapamil, a calcium channel blocker^69^; piribedil, a D2/D3 agonist^70^; luteolin kaempferol and quercetin, flavonoids with potential anti-dementia effects^71^; harmine, an alkaloid which may inhibit tau phosphorylation in AD^72^ etc. Interestingly, quite a number of NSAIDs appeared on our top list of drugs, such as meclofenamic acid, ketorolac, celecoxib, naproxen and acemetacin. NSAIDs have been tested in clinical studies for possible prevention or treatment of AD, although the results were inconsistent and further investigations are required^73^.

#### Attention deficit hyperactivity disorder

A few drugs on the top list have been tested in clinical trials. The anticonvulsant carbamazepine was analyzed in a meta-analysis which concluded preliminary evidence in treating ADHD. The alkaloid lobeline may improve working memory in adult ADHD^74^. Tranylcypromine, another MAOI among the top list, was effective in clinical trials although its side effects need to be considered^75^. Other drugs such as rolipram and reserpine were found to be effective for ADHD-like symptoms in animal models^76,77^.

#### Autistic spectrum disorders

A few drugs on the top list are worth mentioning. Risperidone is one of the two FDA-approved medication for treating irritability in ASD. Two drugs, pentoxylline (phosphodiesterase inhibitor) and amantadine (NMDA antagonist), have been tested in clinical trials of ASD as combination treatment with risperidone. Both showed effectiveness in ameliorating behavioral problems^78,79^. Another drug loxapine, a typical anti-psychotic, was effectiveness as an add-on therapy for irritability in ASD^80^. Ribavirin is another interesting candidate. Recently autism was shown to be associated with eIF4E overexpression which in turn leads to excessive translation of neuroligins^81^. Ribavarin, an antiviral agent, was found to be an inhibitor of eIF4E^82^ and hence may serve as a potential treatment. Ribavarin was also listed in a patent for autism treatment (www.google.ch/patents/US5008251).

### Drug-set enrichment analyses

#### Enrichment for drugs listed in the ATC classification system

First we considered the enrichment test results from drugs listed in the ATC classification system. As shown in Table 4, antipsychotics were strongly enriched in the repositioning candidates for SCZ (lowest *p* across four tests = 4.69E-06) and BD (lowest *p* = 2.26E-07). Interestingly, antipsychotics were also enriched in the drug candidates (albeit less strongly) for MDD (lowest p = 0.0285), AD (lowest p = 0.0256) and anxiety disorders (lowest p =0054). We also observed antidepressants and anxiolytics to be enriched in drugs repositioned for bipolar disorder (lowest p = 1.17E-05). In addition we found a trend towards significance for AD (lowest p = 0.0507). When all psychiatric medications are combined as a drug-set, evidence of enrichment was found for SCZ, BD, AD, and ANX.

**Table 4.**
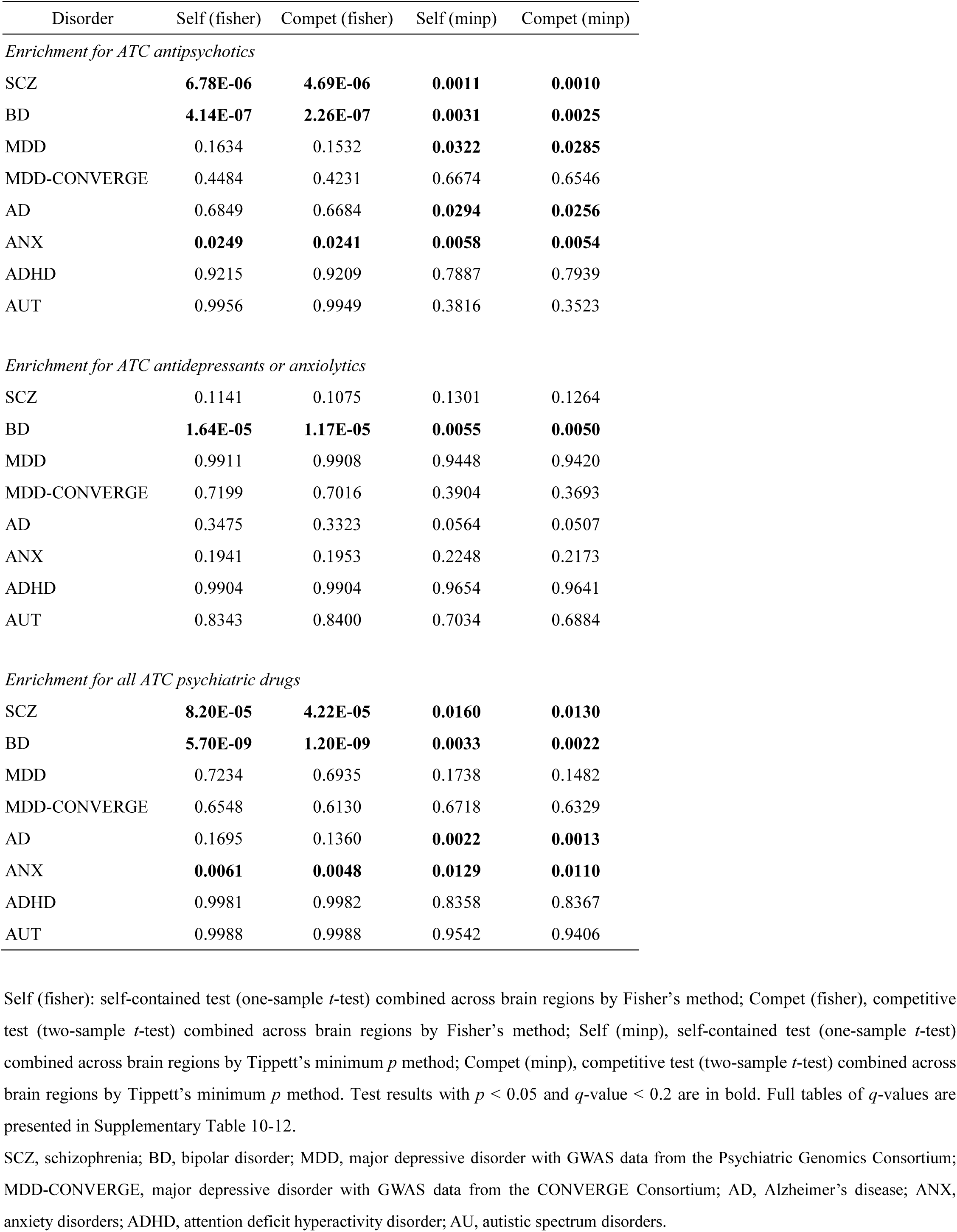
Drug enrichment analyses with drug-sets defined in the Anatomical Therapeutic Chemical (ATC) Classification System

#### Enrichment for drugs listed in MEDI-HPS

The enrichment test results using MEDI-HPS drugs were in general consistent with those derived from the ATC drugs (Table 5). Again we observed that antipsychotics were highly enriched in the repositioning candidates for schizophrenia (lowest p = 2.24E-09). For the rest of the analyses with the antipsychotics drug-set, we found enrichment for BD and MDD; a nominally significant result was also observed for AD.

**Table 5.**
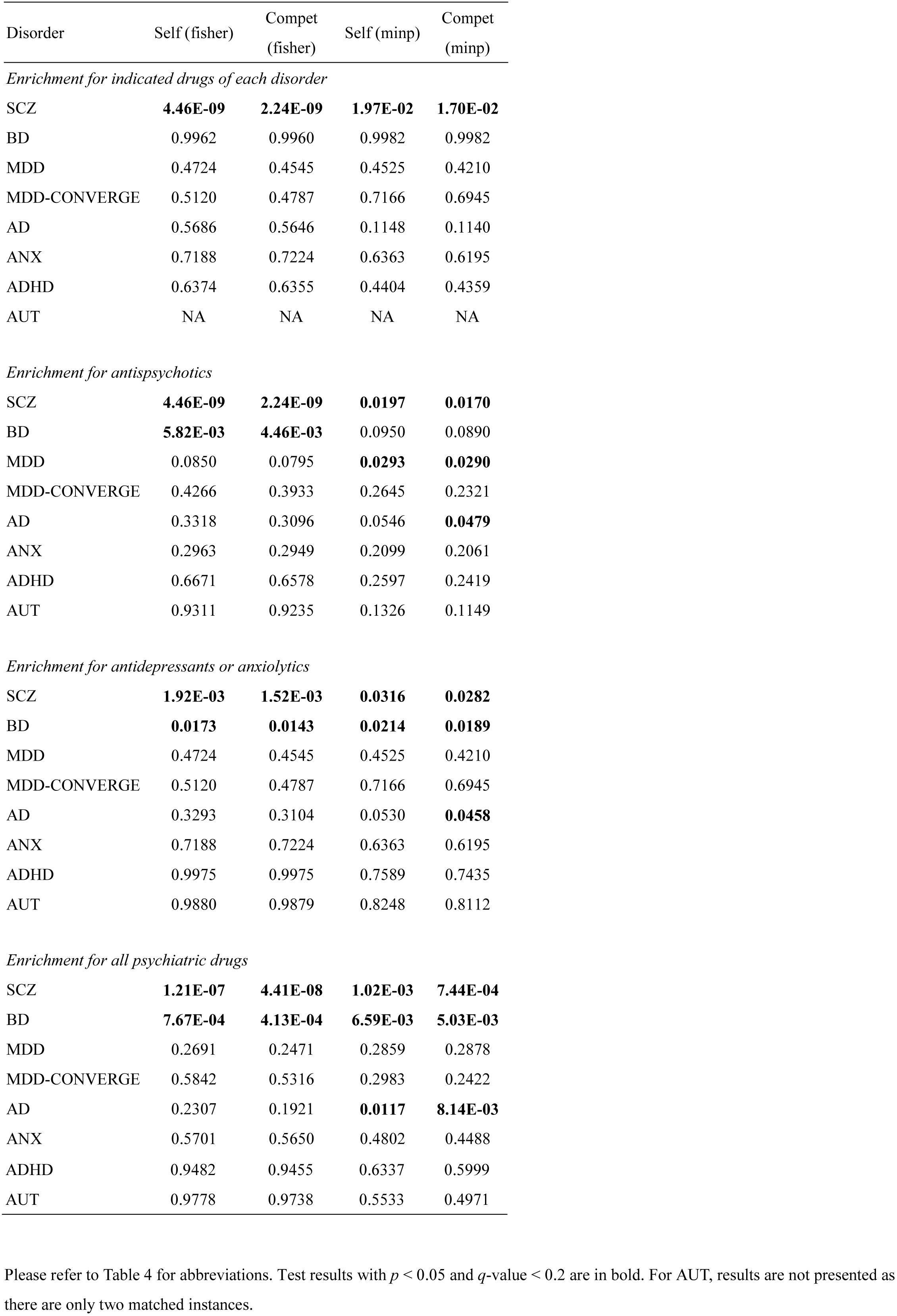
Drug enrichment analyses with drug-sets defined by MEDI-HPS

When the drug-set was limited to antidepressants and anxiolytics, enrichment was observed for SCZ (lowest p = 1.52E-03), BD (lowest p = 0.0143) and AD (lowest p = 0.0458). When all psychiatric drugs were included, enrichment was found for SCZ, BD and AD.

#### Enrichment for drugs listed in clinicalTrial.gov

We tested for enrichment for drugs listed in clinicalTrial.gov for each of the corresponding disorders (Table 6). (We did not pursue tests of drugs across diagnoses as drugs listed in clinicalTrial.gov are less certain and well-defined as a “drug class” compared to the previous two sources). Evidence of enrichment was observed for SCZ (lowest p = 0.0116), BD (lowest p =0.0132), MDD (lowest p = 0.0396) and anxiety disorders (lowest p = 0.0066).

**Table 6.** Drug enrichment analyses with drug-sets defined by those listed in clinicalTrial.gov

| Disorder | Self<br>(fisher) | Compet<br>(fisher) | Self<br>(minp) | Compet<br>(minp) |
| --- | --- | --- | --- | --- |
| SCZ | **0.0162** | **0.0116** | 0.5155 | 0.4949 |
| BD | **0.0167** | **0.0132** | **0.0178** | **0.0158** |
| MDD | **0.0448** | **0.0396** | 0.1085 | 0.1068 |
| MDD-CONVERGE | 0.5465 | 0.4978 | 0.6540 | 0.5910 |
| AD | 0.4371 | 0.4159 | 0.4969 | 0.4642 |
| ANX | 0.0894 | 0.0783 | **0.0100** | **0.0066** |
| ADHD | 0.1765 | 0.1728 | 0.2834 | 0.2691 |
| AUT | 0.9502 | 0.9555 | 0.8904 | 0.9004 |

Please refer to Table 4 for abbreviations. Test results with $p < 0.05$ and $q\text{-value} < 0.2$ are in bold.

## DISCUSSION

In this study we developed a novel framework for drug repositioning by linking two apparently disparate sources of information, GWAS and the Connectivity Map. We proposed to compare the GWAS-imputed transcriptome profiles with those derived from drugs. We applied the methodology to 7 psychiatric disorders and identified a number of interesting candidates for repositioning, many of which are supported by animal studies or clinical trials. The drug-set enrichment analyses further lend support to the validity of our approach.

There are a number of advantages of our repositioning framework. Firstly this approach is largely “hypothesis-free” in the manner that it does not assume any knowledge about known drug targets or drug-disease relationships. As a result the method may be able to find drugs of different mechanisms from the known treatments. This method may be particularly useful for diseases with few known treatments or if existing treatments are highly similar in their mechanisms. A related advantage is that it can be applied to any chemicals as long as the expression profile is available, such as traditional Chinese medicine (TCM) products with no known drug targets^83^. As described before, another important advantage when compared to conventional connectivity mapping is that GWAS-imputed transcriptome is relatively immune to confounding by medication effects. With the availability of eQTL reference resources such as GTEx, imputation of expression profiles can be performed easily for close to a hundred tissues. In addition, only GWAS summary statistics are required as input for imputation, which obviates the difficulties in obtaining raw genotype data and makes the approach easily applicable to a wide variety of traits. The current method is also intuitive and computationally simple to implement. Moreover, we have considered all genetic variants instead of just the most significant hits in our repurposing pipeline. One alternative for inclusion of sub-threshold associations is to employ gene-set analysis to look for over-representation of genes acting as drug targets^84^, but here we proposed a different novel approach that does not rely on knowledge of known drug targets and takes into account the direction of genetic associations.

It is encouraging to observe that our repositioning results are in general supported by the drug-set enrichment analyses. In particular, antipsychotics, which are known to treat SCZ and BD, are also strongly enriched in the repositioning candidates of these two disorders. Similarly, antidepressant is an indicated treatment option of BD^49^, which was also significantly enriched for this disorder. For MDD (using the PGC data) and anxiety disorders, while we did not observe enrichment in ATC or MEDI-HPS drug-sets, there was some evidence that our approach preferentially picked up drugs included in clinical trials. These results suggest that GWAS results contain useful information for drug discovery or repositioning, and is concordant with the conclusion of a previous study that drugs supported by human genomic data increase along the development pipeline^7^.

Interestingly, our analyses also revealed possible enrichment of antipsychotics for MDD, anxiety disorders and AD. Antipsychotics have long been used for treatment of depression, especially in the more severe cases with psychotic symptoms. A recent meta-analysis also demonstrated efficacy of a number of atypical antipsychotics as adjunctive treatment in depression^85^. Given the high co-morbidity of depression and anxiety^86^ and possibly shared pathophysiology^87^, it is not unexpected that antipsychotics may be useful for anxiety as well. Anti-psychotics are not infrequently prescribed for anxiety disorders, although its safety and efficacy warrant further investigations^88^. In a similar vein, we observed that antidepressants and anti-anxiety agents were over-represented in the top drug lists of SCZ and AD (with nominally significance for AD). Psychotic and depressive symptoms are very commonly seen^89^ in AD patients, hence the enrichment for antipsychotics and antidepressants are expected. Antidepressants have been tested in clinical trials for schizophrenia, especially for negative symptoms, although further studies are required in this area^90^. Our findings provide a genetic basis for the effectiveness of psychiatric drugs across diagnoses.

We observed that for some of the psychiatric disorders there were no significant enrichment of known drug indications, or the enrichment came from another class of drugs. There are a few possible explanations. Firstly, the sample size may not be large enough to detect enrichment of known drugs. Among the eight datasets, the sample size of SCZ GWAS is the largest, reaching almost 80000. For the other traits, the sample sizes are mostly between 5000 to 20000. Limited sample sizes imply that some true associations are not detected and the imputed transcriptome is less accurate, which will affect the ability to find drugs with matched (reversed) expression profiles. For any high-throughput studies with limited sample sizes, it is possible for some signals (here known drug indications) to be “missed”, although other associations may be detected. An analogy is that earlier SCZ GWAS with smaller sample sizes did not detect the *DRD2* locus but revealed other loci such as *ZNF804A^91^*. With accumulation of samples, the latest GWAS meta-analysis did confirm *DRD2* as a susceptibility gene^23^. Hence limited sample sizes may explain, for instance, the enrichment of antipsychotics for MDD but not antidepressants.

Secondly, many psychiatric disorders are known to be heterogeneous. For MDD, anxiety disorders and AD, the prevalences are relatively high^92,93^ and there may be a wider range of heterogeneity in clinical manifestations and the underlying pathophysiologies. The heterogeneity impairs study power and implies that a specific drug or drug class may only be effective for a certain group of patients. Third, there are limited available treatment for some disorders, especially AD, AUT and ADHD, which makes the detection of these known drug indications difficult.

In addition to the above, we observed that enrichment analyses were mostly negative for the MDD-CONVERGE dataset. One possible explanation is that as the GTEx dataset from which transcriptome imputation is based is not well-matched to the MDD-CONVERGE sample. The latter sample is composed of Chinese women, while the GTEx project includes only 1.0% Asians and 34.4% of females to date (http://www.gtexportal.org/home/tissueSummaryPage, accessed 22^nd^ Dec 2016). While we expect overlap in the genetics of expression regulation, the imputation quality may be affected. Notwithstanding some negative results in the drug enrichment analyses, many of the top candidates for repositioning are suggested in preclinical or clinical studies, and are still worthy of further investigations.

There are a few general limitations to our approach. It is worth noting that the GWAS-imputed transcriptome captures the genetically regulated part of expression, and expression changes due to other factors (*e.g.* environmental risk factors) cannot be modelled. The method of comparing expressions is largely “hypothesis-free” as mentioned previously, however it may be improved by incorporating knowledge on drug targets or drug-disease relationships, if such information is available. The drug expression profiles were not measured in brain tissues in Cmap, although the original publication on Cmap showed that drugs on neuropsychiatric diseases such as Alzheimer’s disease or schizophrenia can still be reasonably modelled^13^. It should be emphasized that the best approach to verify the repositioning predictions should rest on careful and adequately-sized preclinical and clinical studies, and the current study does not provide confirmatory evidence for the repositioning candidates.

In conclusion, we have developed a novel framework for drug repositioning by linking up GWAS and drug expression profiling and applied the methodology to 7 psychiatric disorders. Our analyses also provide support that the psychiatric GWAS signals are enriched for known drug indications. We also present a list of repositioning candidates for each disorder, which will we believe will serve as a useful resource for preclinical and clinical researchers to pursue further studies.

## Acknowledgements

This study is partially supported by the Lo-Kwee Seong Biomedical Research Fund and a Direct Grant from the Chinese University of Hong Kong. We would like to thank Mr. CHAN Chun Wing for assistance in the annotation of drugs. We are grateful to Professor Stephen K.W. Tsui for useful discussions and the Hong Kong Bioinformatics Centre for computing support.

